# Quantitative Insights into Processivity of an Hsp100 Protein Disaggregase on Folded Protein Substrates

**DOI:** 10.1101/2024.10.09.617403

**Authors:** Jaskamaljot Kaur Banwait, Aaron L. Lucius

**Affiliations:** Department of Chemistry, University of Alabama at Birmingham, Birmingham, Alabama

## Abstract

The Hsp100 family of proteins play important roles in maintaining protein homeostasis in cells. *E. coli* ClpB is an Hsp100 protein that remodels misfolded proteins or aggregates. ClpB is proposed to couple the energy from ATP binding and hydrolysis to processively unfold and translocate protein substrates through its axial channel in the hexameric ring structure. However, many of the details of this reaction remain obscure. We have recently developed a transient state kinetics approach to study ClpB catalyzed protein unfolding and translocation. In this work we have used this approach to begin to examine how ATP is coupled to the protein unfolding reaction. Here we show that at saturating [ATP], ClpB induces the cooperative unfolding of a complete TitinI27 domain of 98 amino acids, which is represented by the kinetic step-size m ∼100 amino acids. This unfolding event is followed by rapid and undetected translocation up to the next folded domain. At sub-saturating [ATP], ClpB still induces cooperative unfolding of a complete TitinI27 domain but translocation becomes partially rate-limiting, which leads to an apparent reduced kinetic step-size as small as ∼ 50 amino acids. Further, we show that ClpB exhibits an unfolding processivity of P = (0.74 ± 0.06) independent of [ATP]. These findings advance our understanding of the elementary reactions catalyzed by E. coli ClpB but are broadly applicable to a variety of Hsp100 family members.

**Graphical Abstract:** 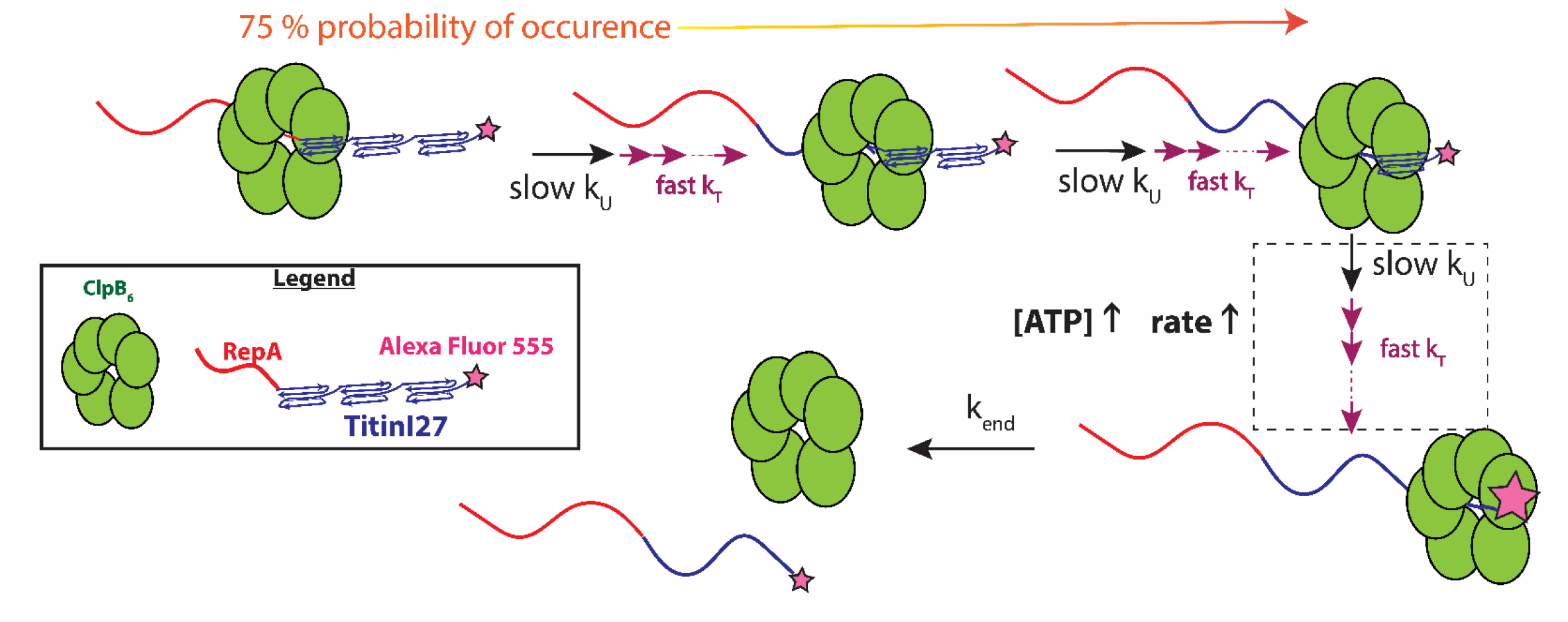

## Introduction

ATPases associated with diverse cellular activities (AAA+) are essential for cellular functions such as organelle biogenesis, RNA processing, membrane fusion, DNA replication, repair, recombination, and chromatin remodeling [1-5]. Across domains of life, representative AAA+ molecular motors are essential for proteome maintenance [6]. *E. coli* ClpB and *S. cerevisiae* Hsp104 are AAA+ protein disaggregases that, in collaboration with the co-chaperones, DnaK, DnaJ, and GrpE (KJE), resolve protein aggregates that form during heat shock or stress [7-12]. Interestingly, metazoans lack a ClpB/Hsp104 homologue.

The molecular mechanisms of protein disaggregation and collaboration with co-chaperones are not fully understood. ClpB is proposed to couple ATP binding and hydrolysis to processive rounds of protein unfolding and translocation of the newly extracted polypeptide chain through the axial channel of the hexameric ring structure of the enzyme.

ClpB contains two ATP binding and hydrolysis sites per monomer termed nucleotide binding domain (NBD) 1 and NBD 2. Like most AAA+ molecular motors, ClpB requires nucleoside triphosphate binding to drive the assembly of a hexameric ring with polypeptide binding activity [13-15]. Assembly leads to twelve ATP binding and hydrolysis sites per active hexameric ring. Interestingly, in the case of both ClpB and Hsp104, only the slowly hydrolysable nucleotide analogue, ATPγS, will drive the formation of hexamers with polypeptide binding activity. However, neither ATP nor the non-hydrolysable nucleotide analogues, AMP-PNP and AMP-PCP, support the formation of hexamers with polypeptide binding activity [16].

Wickner and coworkers discovered that a 1:1 mixture of ATP:ATPγS could activate ClpB in a manner that alleviated the need to include KJE [17]. This strategy has been exploited to examine the activities of ClpB in the absence of the complicating factors introduced by adding three additional proteins/enzymes to an *in vitro* experiment. Although it is not fully understood how ATPγS activates ClpB, the initial explanation for this phenomenon was that slow hydrolysis of ATPγS at a subset of the twelve ATP binding/hydrolysis sites in ClpB is needed to activate the enzyme. The dominant explanation that has emerged since that work is that ATPγS binds to the D1 ATPase sites, activates the motor, and ATP hydrolysis at D2 drives rounds of ClpB catalyzed protein unfolding and translocation[18].

Most recently, we have shown that ATPγS alone can support processive rounds of protein unfolding catalyzed by ClpB [19]. This observation led us to develop a sequential mixing stopped-flow strategy that controls for ATPγS driven translocation and reports on processive rounds of ClpB catalyzed protein unfolding. In this design, the first mixing event allows for ATPγS driven hexamer formation followed by binding to a model soluble folded protein targeted to be unfolded. The second mixing event rapidly adds hydrolysable ATP. From that work we showed that ClpB processively unfolded up to three tandem repeats of the Titin I27 domain with an overall rate of ∼0.9 and ∼3 amino acids per second in the presence of 500 μM or 150 μM ATPγS, respectively, and a fixed concentration of 500 μM ATP. Additionally, we reported a kinetic step-size of ∼60 amino acids unfolded between two rate limiting steps.

The Titin I27 domain has been shown to cooperatively unfold in both atomic force microscopy (AFM) and optical tweezer experiments [20, 21]. Thus, it seems unlikely that the observed kinetic step-size of ∼60 amino acids represents a mechanical unfolding step-size. Rather, we hypothesize that ClpB catalyzes the complete unfolding of a 98 amino acid Titin I27 domain and rapidly translocates on the newly unfolded polypeptide chain up to the next folded Titin I27 domain. Protein unfolding is rate-limiting but translocation is partially rate-limiting at the sub-saturating ATP concentrations of 500 μM ATP reported by us [22]. Consequently, the kinetic step-size underestimates the expected mechanical unfolding step-size of 98 amino acids.

To test the hypothesis that unfolding is rate-limiting we examined the complete ATP concentration dependence of the kinetic parameters. Here we show that ClpB catalyzes protein unfolding with a kinetic step-size of ∼100 amino acids between two rate-limiting steps and a maximum rate of ∼30 amino acids s^-1^ at saturating ATP concentrations and a fixed 150 μM ATPγS. The consistency between the kinetic step-size and the length of a 98 amino acid Titin I27 repeat suggests that ClpB catalyzes the cooperative unfolding of a complete Titin I27 domain. Thus, protein unfolding is fully rate-limiting at saturating ATP, and translocation is fast and undetected. However, at limiting [ATP], translocation becomes partially rate-limiting leading to a kinetic step-size that is smaller than the mechanical unfolding step-size. Further, we show that ClpB exhibits a protein unfolding processivity, P, of ∼0.7, independent of [ATP]. Finally, we show that the rate depends cooperatively on [ATP] consistent with cooperative communication between the twelve ATP binding and hydrolysis sites during enzyme catalyzed protein unfolding in this AAA+ hexameric ring motor.

## Results

### ATP Concentration Dependence of ClpB Catalyzed Protein Unfolding

To interrogate the protein unfolding cycle as a function of [ATP] we performed sequential mixing stopped-flow experiments as schematized in Fig. 1 A using the three RepA-Titin_X_ constructs illustrated in Fig. 1 B and previously introduced in [19]. In this experiment, 8 μM ClpB is loaded into syringe 1 and rapidly mixed with the contents of Syringe 2, containing 600 μM ATPγS and 200 nM RepA-Titin_X_, where X = 1, 2, or 3 repeats of the Titin I27 domain. Upon rapid mixing the sample is held in the ageing loop for a user-defined amount of time, Δt_1_ = 10 minutes. The elapsed time, Δt_1_, allows ClpB to bind ATPγS and assemble into hexameric rings active in binding to the unstructured N-terminal RepA sequence on the substrates. Further, using the energy from the slow hydrolysis of ATPγS, ClpB begins to slowly unfold and translocate the polypeptide chain. After Δt_1_ = 10 minutes elapses, the sample is rapidly mixed with the contents of Syringe 3, which contains varying amounts of ATP and 40 μM α-casein to serve as a trap for free ClpB. The sample flows into the observation chamber and fluorescence from Alexa Fluor (AF) 555 at the C-terminus of the substrate is monitored. All mixing events are 1:1, which leads to a final reaction concentration of 2 μM ClpB monomer, 150 μM ATPγS, 50 nM RepA-Titin_X_, 20 μM α-casein, and the ATP concentrations indicated.

**Figure 1.**
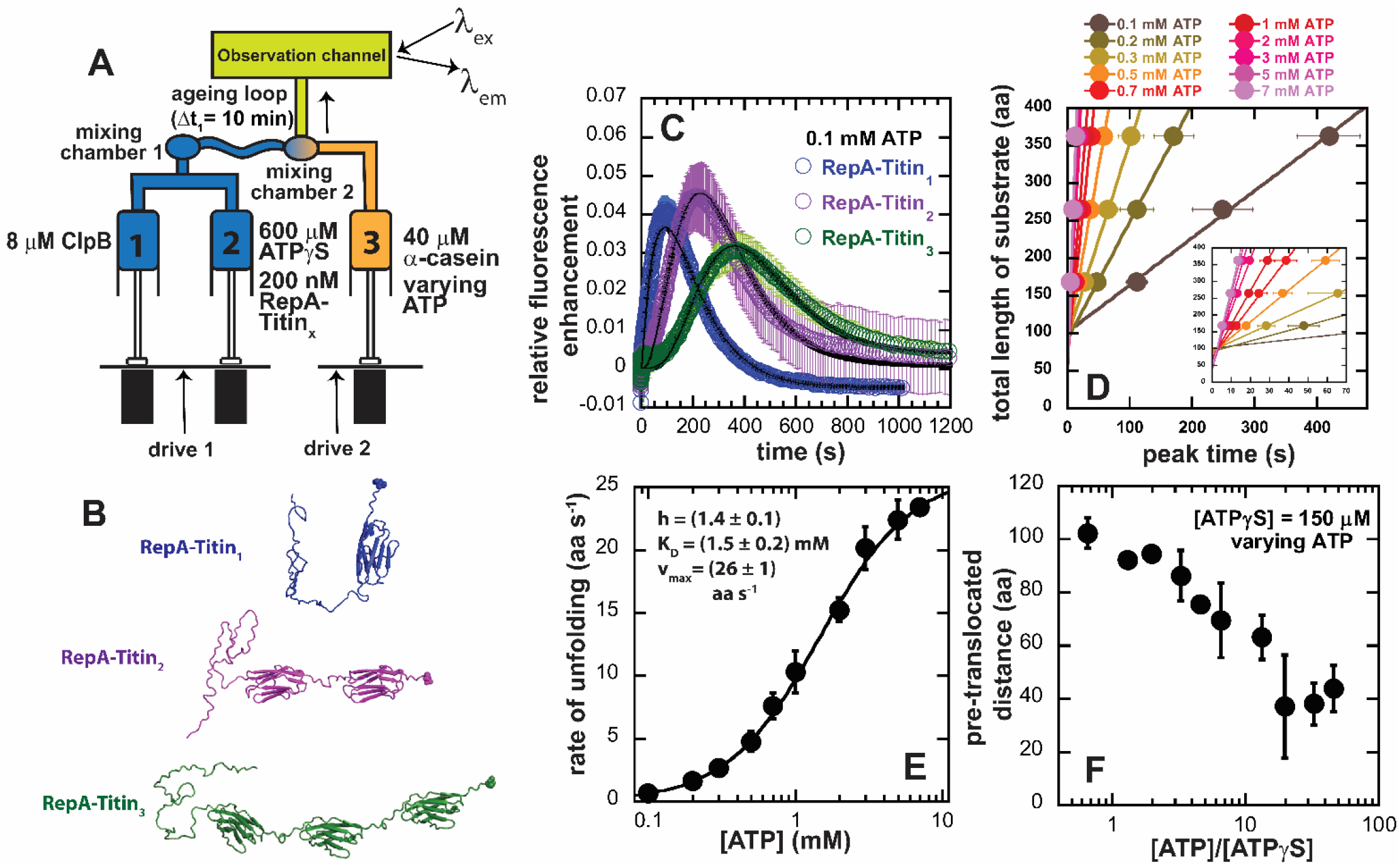
Varying [ATP] to investigate ATP coupled unfoldase activity of ClpB A) Sequential-mix stopped flow schematic for [ATP] dependence: Single turnover stopped-flow experiments are carried out by varying ATP in syringe 3. Syringe 1 contains 8 mM ClpB. Syringe 2 contains 600 μM ATPγS and 200 nM RepA-Titin_X_. The components of syringes 1 & 2 are rapidly mixed in mixing chamber 1 and then they age in the aging loop for a fixed Δt_1_ of 600 s. We expect hexameric ClpB to pre-bind RepA-Titin_X_ during Δt_1_. Syringe 3 contains 40 μM α-casein and varying [ATP]. After Δt_1_ has elapsed the pre-bound complex is rapidly mixed with components of syringe 3 in mixing chamber 2. From there, the reagents travel to the observation channel where Alexa Fluor 555 is excited at 520 nm and the emission signal is monitored at λ_em_ ≥ 570 nm. B) Structures of RepA-Titin_1_ (blue), RepA-Titin_2_ (purple), and RepA-Titin_3_ (green). Each substrate has unstructured RepA (1-70) on the N-terminus followed by (TitinI27)_X_, where x denotes the number of folded β sandwich structures of TitinI27. Each substrate has a Cysteine on the C-terminus represented by a space-filling model. This Cysteine is covalently attached with Alexa Fluor 555 dye via Cysteine maleimide reaction (not shown). C) Representative time courses collected for RepA-Titin_1_ (blue), RepA-Titin_2_ (purple), and RepA-Titin_3_ (green) using schematic A at 100 μM ATP. Each time course represents an average of three or more sequential time courses and error bars represent the standard deviation of the averaged time courses. The solid black lines represent the best fit from fitting the data to scheme 1 in Fig. 2 A. The fitting parameters obtained are *m* = (61 ± 4) aa and *k*_*u*_ = (0.130 ± 0.004) s^-1^. D) Plot of the total length of substrate as a function of peak time determined for 0.1 mM, 0.2 mM, 0.3 mM, 0.5 mm, 0.7 mM, 1 mM, 2 mM, 3 mM, 5 mM and 7 mM of ATP. The inset plot shows the first 70 seconds of peak time to display data from the highest concentrations of ATP being investigated. The solid lines represent linear fits yielding E) slopes versus [ATP] F) intercepts versus [ATP]/ [ATPγS]. All data points and error bars represent the average and standard deviation determined from three replicates. E) rate of unfolding plotted as a function of [ATP] is fit to Hill equation, eq 1. The solid line represents the best fit line and the parameters obtained are *h* = (1.4 ± 0.1), *K*_*D*_ = (1.5 ± 0.2) mM and *V*_*max*_ = (26 ± 1) aa s ^-1^.

Example time courses for experiments carried out with the three different substrates shown in Fig. 1 B are shown in Fig. 1 C for a final [ATP] = 0.1 mM. Each time course exhibits a lag phase followed by fluorescence enhancement, which we have shown indicates PIFE due to the arrival of ClpB at the C-terminal fluorophore, AF555 [19]. The maximum PIFE signal is followed by loss of fluorescence due to dissociation of ClpB. A length-dependent series of time courses were collected for final ATP concentrations varying between 0.1 mM and 7 mM in the presence of 150 μM ATPγS.

As previously reported, the time at which the peak in fluorescence occurs increases with increasing number of Titin I27 domains or total substrate length, *L*. Thus, for each [ATP], the total length was plotted vs. the observed peak time. In principle, this results in a classical kinematics position vs. time plot, see Fig. 1 D and inset for high [ATP].

For each [ATP] the plot was observed to linearly increase and exhibit a slope that represents the rate of ClpB catalyzed protein unfolding. This overall rate was plotted vs. [ATP] and is shown in Fig. 1 E. The overall rate of unfolding vs. final [ATP] exhibits cooperativity and is best described by a Hill model, given by Eq. 1, compared to a rectangular hyperbola, see Supp. Fig. 1 for a fit to Eq. 1 with *h* = 1. The rate vs. ATP concentration was subjected to non-linear least squares (NLLS) analysis using Equation 1 and a maximum rate of (26 ± 1) amino acid s^-1^, *K*_*D*_ = (1.5 ± 0.2) mM, and a hill coefficient of *h* = (1.4 ± 0.1) was determined.

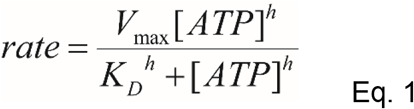

Each plot of total length of substrate vs. peak time exhibits a non-zero substrate length intercept at zero peak time, Δt_2_ = 0. This implies that some length of the substrate has already been unfolded and/or translocated during Δt_1_, i.e. before mixing with ATP. We have previously interpreted this observation to be the result of ATPγS driven translocation that occurs during Δt_1_ and have termed this phenomenon the pre-translocated distance. Here, the pre-translocated distance was observed to decrease with increasing [ATP], see Fig. 1 F. This is surprising because the amount of time that the ClpB-(RepA-Titin_X_) ternary complex is exposed to ATPγS is the same for all final ATP concentrations. Consequently, if this number represents the distance pre-translocated during Δt_1_ then it is expected to be independent of [ATP]. However, any length of the substrate that is not part of the unfolding and translocation reaction, whether at the beginning or end of the reaction, will contribute to the magnitude of the intercept determined from the fit of substrate length vs. peak time in Fig. 1 D. Thus, at low [ATP] ClpB may dissociate before complete translocation of the final Titin I27 domain. Whereas, at high ATP the motor may fully translocate to the end. Although the individual measurements shown in Fig. 1 F are outside of the error of one another, the trend may be an artifact reflecting the reliability of determining the intercept as the slope increases.

### Deconvoluting the overall rate

The peak time analysis presented in Fig. 1 C – F yields an overall rate of protein unfolding, i.e. amino acids per second. However, the overall rate is a convolution of the rate constant defining each repeating step, *k*_*U*_, and the amount of substrate unfolded or translocated between each repeating rate-limiting step, or the step size, *m*. Thus, to determine the kinetic step-size and the elementary stepping rate constant(s) we subjected each set of length-dependent time courses to global analysis using Scheme 1, see Fig. 2 A.

**Figure 2.**
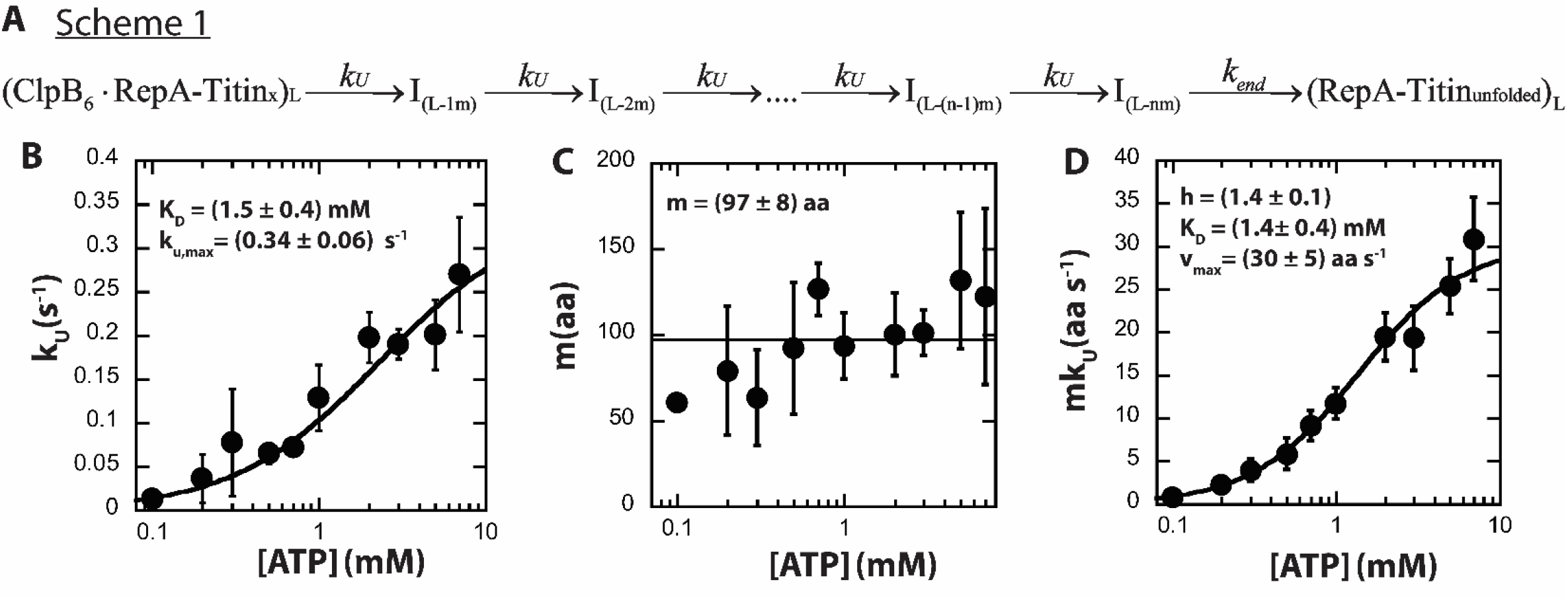
*n*-step sequential mechanism describes ATP coupled unfoldase activity of ClpB: A) Proposed kinetic scheme for ClpB catalyzed protein unfolding and translocation on RepA-Titin_X_ substrate. Parameters B) *k*_*U*_ C) *m* D) *mk*_*U*_ obtained from fitting to scheme 1 at each [ATP] are shown in solid black circles. The black solid line represents the best-fit line. The plot of B) *k*_*U*_ as a function of [ATP] is fit to eq 2 and the parameters obtained are *K*_*D*_ = (1.5 ± 0.4) mM and *k*_*U,max*_=(0.34 ± 0.06) s^-1^. The plot of C) m as a function of [ATP] is fit to a constant at *m* = (97 ± 8) aa. The plot of D) *mk*_*U*_ as a function of [ATP] is fit to eq 1 and the parameters obtained are *h* = (1.4 ± 0.1), *K*_*D*_ = (1.4 ± 0.4) mM, and *V*_*max*_=(30 ± 5) aa s^-1^. All data points and error bars represent the average and standard deviation determined from three replicates.

Scheme 1 starts with hexameric ClpB bound to RepA-Titin_X_ of substrate length, *L*, where the length is indicated by the subscript. A rate-limiting step with rate constant, *k*_*U*_ occurs to form the first intermediate, I, with a substrate length that has been reduced by one step-size, *m*, resulting in a new length of *L* – 1*m*. The reaction then cycles through *n* number of steps before ClpB arrives at the end and slowly dissociates with rate constant *k*_*end*_.

Each set of three time-courses collected at a fixed total [ATP] was subjected to global analysis using Scheme 1, where the unfolding rate constant, *k*_*U*_, and the kinetic step-size, *m*, and the dissociation rate constant, *k*_*end*_, are constrained to be the same for all three time-courses, i.e. global parameters. The resultant parameters from the simultaneous fitting of three length-dependent time-courses are shown in Fig. 2 B – D. The values and the uncertainty bars represent the resultant parameters from the analysis of three independent replicates.

The unfolding rate constant, *k*_*U*_, exhibits a hyperbolic dependence on [ATP] and is well described by a rectangular hyperbola, Eq. 2. From the analysis we found the maximum unfolding rate constant *k*_*U,max*_ = (0.34 ± 0.06) s^-1^ at saturating [ATP] and *K*_*D*_ = (1.5 ± 0.4) mM. Unlike the rate determined from the peak time analysis strategy in Fig. 1 E, we did not detect cooperativity in the rate constant *k*_*U*_ vs. [ATP].

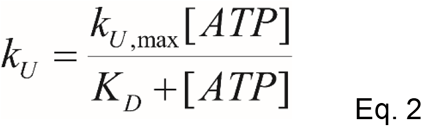

The kinetic step-size, *m*, exhibits a systematic increase with increasing [ATP], see Fig. 2 C. However, the variability in this parameter, between replicates, suggests that all determined values are within error. Assuming that the kinetic step-size is independent of [ATP] we find that the average step-size across all [ATP], <*m*> = (97 ± 8) amino acids. Since each domain of Titin I27 is 98 amino acids, an unfolding step-size of ∼98 amino acids implies that when ClpB applies force to catalyze protein unfolding one domain of Titin I27 cooperatively unfolds in a single rate-limiting step.

From the global analysis, the overall rate is calculated as the product of the kinetic step size, *m*, and the stepping rate constant, *k*_*U*_. The overall rate, *mk*_*U*_, was determined for the three replicates at each [ATP] and are shown in Fig. 2 D. Unlike the rate constant, k_U_ vs. [ATP], the overall rate, mk_U_, depends cooperatively on [ATP]. Thus, the rate vs. [ATP] was subjected to NLLS analysis using the Hill equation given by Eq. 1. From this analysis we determined a Hill coefficient, *h* = (1.4 ± 0.1), a binding constant of *K*_*d*_ = (1.4 ± 0.4) mM, and maximum overall rate of *mk*_*U,max*_ = (30 ± 5) aa s^-1^. Thus, the parameters describing the overall rate determined from peak time analysis and global fitting are in excellent agreement, compare Fig.1 E to Fig. 2 D.

### ClpB concentration dependence of unfolding mechanism

In a single turnover experiment, the peak amplitude of the collected time course is proportional to the extent of binding,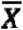, given by Eq. 3. Also, the magnitude of the peak should be proportional to the fraction of ClpB hexamers bound to the substrate scaled by a fluorescence factor, *F*, and the probability that ClpB bound at time zero will fully translocate to the C-terminus to induce PIFE after mixing with ATP. The probability would be given by the processivity, *P*, raised to the power of the number of rate-limiting steps required to reach the end of the substrate, *n*. In sum, the peak amplitude is given by the product of those three parameters as shown in Eq. 4.

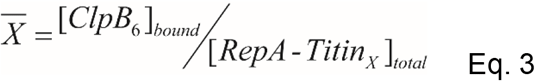

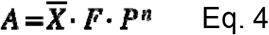

The extent of binding, 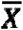, or the amount of hexameric ClpB (ClpB_6_) bound to the RepA sequences would be predicted to be the same in all experiments performed using the mixing strategy illustrated in Fig. 1 A. However, the 200 nM RepA-Titin_X_ used in each experiment is based on the AF555 concentration, see materials and methods. Since the labeling efficiency is not 100 %, each RepA-Titin_X_ sample contains a different fraction of labeled to unlabeled substrate. The unlabeled RepA-Titin_X_ substrate will compete for binding with the labeled substrate but only labeled substrate gives rise to signal. Consistently, RepA-Titin_2_ has the highest labeling efficiency and always exhibits a higher peak amplitude compared to RepA-Titin_1_ and RepA-Titin_3_. This is because the extent of binding to the labeled substrate is higher for RepA-Titin_2_ than for the other two constructs.

In principle, regardless of labeling efficiency, one should be able to saturate the labeled RepA-Titin_X_ with ClpB_6_ at sufficiently high total [ClpB]. We sought to test this by examining different ClpB total concentrations. Sequential mixing experiments were carried out as schematized in Fig. 1 A with a ClpB total concentration of 14, 8, and 4 μM in Syringe 1. After the first mixing event the total concentration drops to 7, 4, and 2 μM ClpB monomer in the presence of 300 μM ATPγS, or 340, 157, and 56 nM ClpB_6_, respectively, based on our previous analysis of the ligand-linked assembly reaction [13, 14, 23].

After Δt_1_ = 10 minutes elapses to allow hexameric ClpB to bind the RepA-Titin_X_ substrate, the bound complex is rapidly mixed with ATP. The resulting time courses collected at a binding concentration of 340 nM ClpB_6_ are shown in Fig. 3 A. The peak amplitudes for each substrate length are shown in Fig.3 B as a function of the predicted hexamer concentration present in the binding reaction. At the highest achievable concentration of ClpB we did observe that the amplitude follows the expected trend of RepA-Titin_1_ > RepA-Titin_2_ > RepA-Titin_3_, see Fig. 3 A and B. The time-courses were subjected to peak time analysis and the rate of unfolding, and the pre-translocated distance were observed to be independent of the ClpB hexamer concentration, see Fig. 3 C - E. This is expected because, under single-turnover conditions, the kinetic parameters should be independent of protein concentration unless there are differentially populated active states. That is to say, if dimers, tetramers, or hexamers of ClpB, that are known to be present [13, 14, 23], bound to the RepA sequence at elevated protein concentration then this would be revealed in the kinetic parameters. Thus, we can conclude that the same active state is bound over this protein concentration range. Moreover, by increasing the [ClpB] we can saturate the extent of binding to the fluorescent substrate.

**Figure 3.**
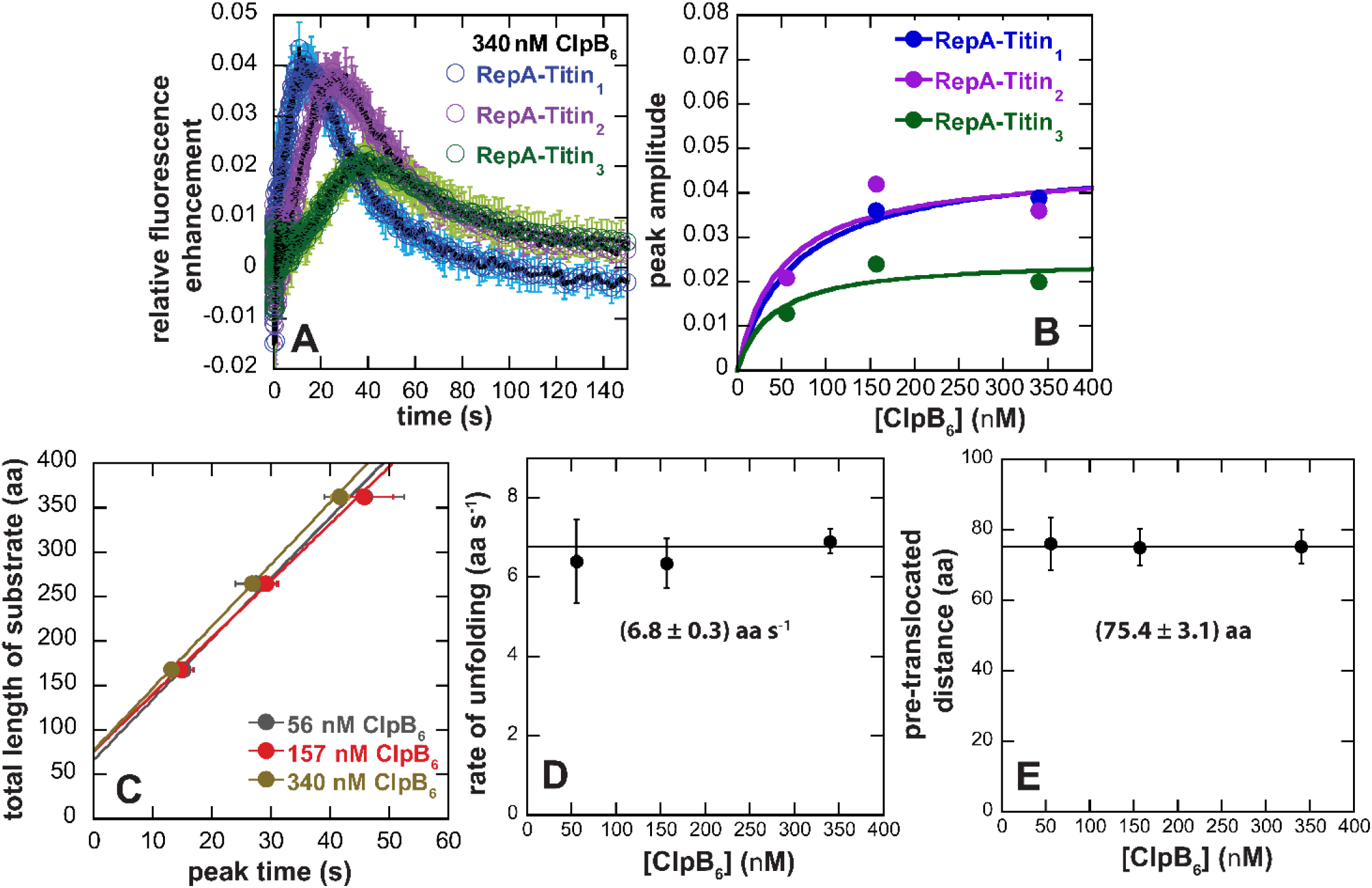
A) Sequential mix stopped flow for [ClpB] dependence: Single turnover stopped-flow experiments are carried out using the same conditions as mentioned in Fig. 1 A except for varying [ClpB] in syringe 1 and fixed 1 mM [ATP] in syringe 3. Representative time courses collected for RepA-Titin_1_ (blue), RepA-Titin_2_ (purple), and RepA-Titin_3_ (green) are shown at 340 mM ClpB_6_ i.e.14 mM ClpB in syringe 1. Each time course represents an average of three or more sequential time courses and error bars represent the standard deviation of the averaged time courses. The solid black lines represent the best fit from fitting the data to scheme 1 in Fig. 2 A. The fitting parameters obtained are *m* = 91(77,96) aa and *k*_*u*_ = (0.12 ± 0.01) s^-1^. B) Peak amplitudes were determined for each substrate at each [ClpB_6_] and plotted here as a function of [ClpB_6_] for RepA-Titin_1_ (blue), RepA-Titin_2_ (purple), and RepA-Titin_3_ (green). The plot was fit to eq 4 and the solid lines represent the best fit. Plot of the total length of substrate as a function of peak time determined for 56 nM (grey), 157 nM (red), and 340 nM (gold) of ClpB_6_. The solid lines represent linear fits yielding D) slopes versus [ClpB_6_] E) intercepts versus [ClpB_6_]. All data points and error bars represent the average and standard deviation determined from three replicates.

### The Peak Amplitude Inversely Depends on Final [ATP]

To interrogate the dependence of the peak amplitude on [ATP], we plotted all the time-courses collected with RepA-Titin_3_ as a function of [ATP], see Fig. 4 A. At 0.1 mM final ATP we observed the highest peak amplitude and, as the concentration of ATP is increased, both the peak magnitude and the signal to noise decreases, see Fig. 4 A and B. This was curious because one expects the signal to improve when providing more energy (ATP) to an ATP-driven motor.

**Figure 4.**
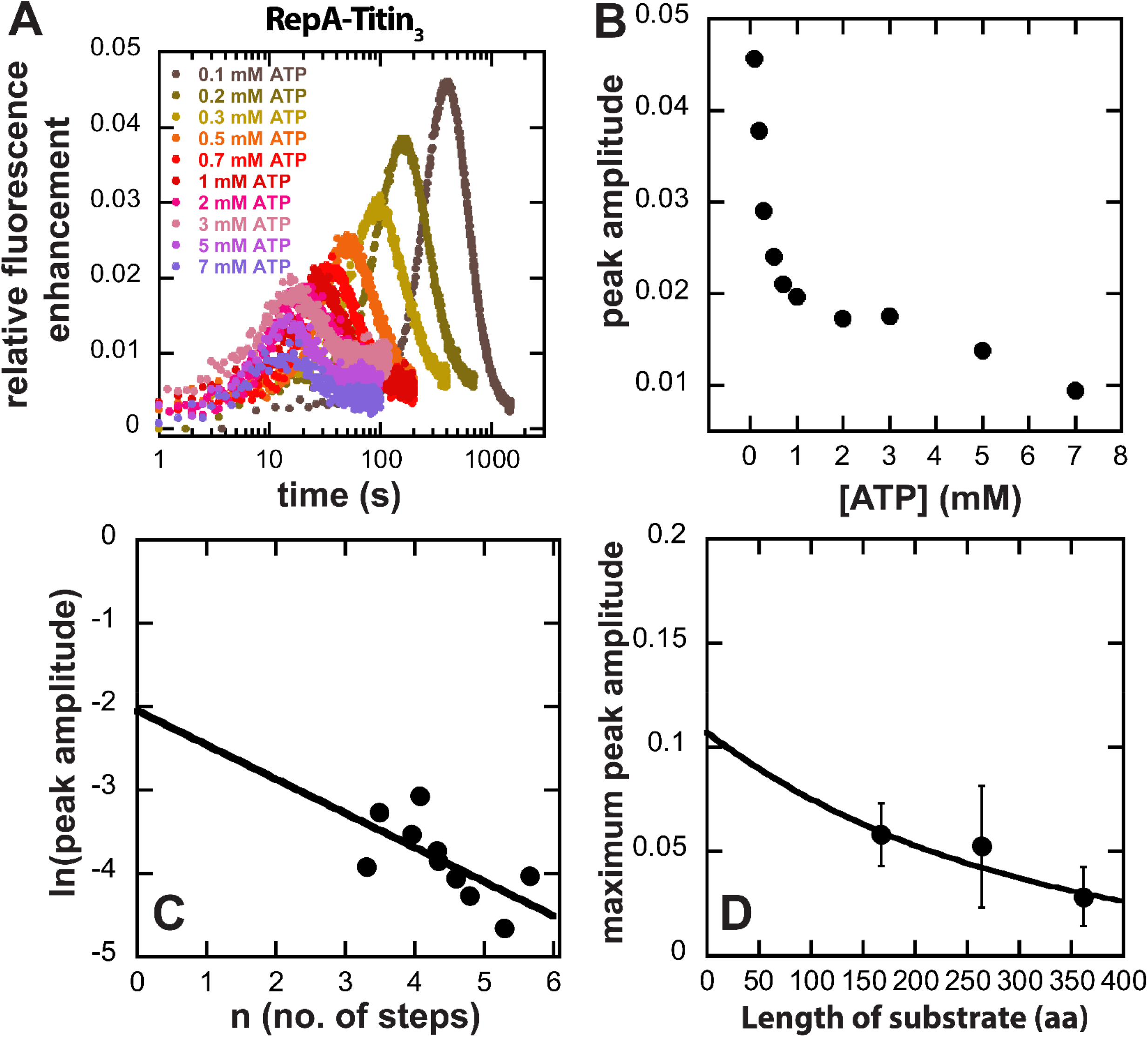
Processivity determination through RepA-Titin_3_ time courses varying with [ATP]: A) Time courses collected for RepA-Titin_3_ using the schematic described in Fig. 2 A at varying [ATP]. As [ATP] increases, peak time shortens. Interestingly, the amplitude at peak time, termed peak amplitude, is also changing with [ATP] which is plotted in (B) as a function of [ATP]. (C) ln of peak amplitude is plotted as a function of n_3_, unique to each [ATP] for RepA-Titin_3_. The plot fits a linear equation 3d represented by a solid black line. Processivity calculated from this analysis is (0.74 ± 0.06). (D) The maximum peak amplitude, calculated by fitting Fig. 3 B to eq 4 for each substrate, was plotted as a function of the total length of the substrate. The black solid line represents the best fit to eq 5 which results in processivity = (0.75 ± 0.17).

**Table 1:** This table summarizes the parameters obtained by fitting Figs. 2 D and 3 D to eq 1.

| **Parameters** | **Fig. 2 D** | **Fig. 3 D** |
| --- | --- | --- |
| **h** | **$1.4 \pm 0.1$** | **$1.4 \pm 0.1$** |
| **K<sub>D</sub>(mM)** | **$1.5 \pm 0.2$** | **$1.4 \pm 0.4$** |
| **v<sub>max</sub>(aa s<sup>-1</sup>)** | **$26 \pm 1$** | **$30 \pm 5$** |

To further analyze this phenomenon, we plotted the peak amplitude vs. total [ATP], see Fig. 4 B. Interestingly, the amplitude precipitously decreases from 0.1 mM ATP to ∼1 mM ATP before an apparent plateau, followed by another decrease between ∼3 and 7 mM ATP. However, the second apparent decrease may be the result of the low signal to noise above 3 mM ATP, see Fig. 4 A. The midpoint of the transition occurs at ∼ 270 μM, which may represent an apparent equilibrium constant for ATP competing with ATPγS.

The reduction in amplitude with increasing [ATP] cannot be explained by differences in labeling efficiency. In this comparison, the labeling efficiency is the same in all experiments since we are only comparing time-courses collected with RepA-Titin_3_. Consequently, the extent of binding, 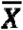(Eqs. 3 and 4), before mixing with ATP, is expected to be the same for each time-course. This is because, after mixing 8 μM total ClpB monomer with 200 nM RepA-Titin_3_, 600 μM ATPγS, and allowing assembly and binding to proceed for ten minutes, 340 nM ClpB_6_ is available to bind to 100 nM RepA-Titin_3_ in the presence of 300 μM ATPγS but no ATP.

We expect that the PIFE signal, *F* in Eq. 4, would not depend on [ATP], i.e. all ClpB that arrive at the C-terminus impact AF555 in the same way regardless of [ATP], especially on the same substrate. Thus, the simplest interpretation of the reduction in amplitude as a function of increasing [ATP] is that less ClpB is arriving at the C-terminal AF555 as the [ATP] is increased. This either means that processivity, *P*, is reduced with increasing [ATP] or, upon rapid mixing with ATP, some fraction of ClpB is induced to dissociate. That is to say, 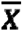 is instantaneously perturbed upon rapid mixing with ATP, or some fraction of bound ClpB_6_ never initiates unfolding and translocation.

To separate the potential impact of extent of binding, 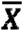, from processivity, P, we turned to Eq. 4. If one executes the natural log of both sides of this equation, then Eq. 4 can be simplified to Eq. 5. Equation 5 predicts that a plot of the natural log of the amplitude, *A* vs. the number of steps, *n*, would yield a linear plot with a slope of the natural log of the processivity, ln(*P*), and an intercept of the natural log of the product of extent of binding, 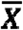, and fluorescence, *F*, ln 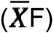.

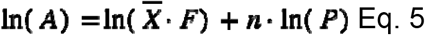

Figure 4 C shows a plot of the natural log of the amplitude, ln(*A*), vs the number of steps from fitting of the time courses in Fig 4 A. The solid black line represents a linear fit with slope = ln(*P*) = -0.41 and an intercept of ln(*A*) = -2.05. The intercept represents the product of extent of binding and fluorescence at zero steps, which represents the fluorescence and extent of binding at the instance of the second mixing event, i.e. addition of ATP. By exponentiating the slope we determine a processivity, *P* = 0.7 ± 0.1, which assumes that the processivity is independent of [ATP]. If the processivity is independent of [ATP] then this observation reveals that upon rapid mixing with ATP some fraction of bound ClpB immediately dissociates thereby perturbing 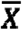. But all molecules that initiate protein unfolding and translocation exhibit the same processivity of ∼ 0.7.

As equation 4 shows, the peak amplitude, *A*, is proportional to the product of the extent of binding, fluorescence, and processivity. If one assumes that the extent of binding for all three RepA-Titin_X_ substrates is the same at the highest ClpB concentration tested (ClpB_6_ = 340 nM), see Fig. 3 B, and the impact on fluorescence, F, due to ClpB arriving at the fluorophore is the same for each substrate length then a plot of peak amplitude vs. substrate length will be proportional to the processivity, *P*, raised to the power of the number of steps, *n*. To test this the peak amplitude from the time courses in Fig. 3 A plotted vs. substrate length, *L*, and is shown in Fig. 4 D [24, 25]. The solid line represents the result of a NLLS fit using Eq. 4. From this analysis a processivity of P = 0.7 ± 0.2 was determined, which is in good agreement with *P* = 0.7 ± 0.1 determined from ln(*A*) vs. *n* in Fig. 4 C, under the assumption that the processivity is independent of [ATP].

The agreement between the processivity determined when assuming *P* is independent of [ATP] or assuming that extent of binding is independent of length suggests that processivity is the same for all hexamers bound that initiate protein unfolding. Thus, we propose that the most likely explanation for the reduction in peak amplitude with increasing [ATP] is that, upon mixing with increasing concentrations of ATP, some fraction of hexamers dissociate or do not initiate translocation.

## Discussion

*E. coli* ClpB and *S. cerevisiae* Hsp104, in collaboration with co-chaperones, catalyze the reversal of protein aggregates that form during heat shock or stress. Specifically, ClpB collaborates with the co-chaperones DnaK, DnaJ, and GrpE (KJE) to reverse protein aggregates. Over thirty years of work has been done on this system, and a great deal has been learned [26, 27]. However, barriers remain in place to a complete and quantitative understanding of the molecular mechanisms.

ClpB has long been hypothesized to catalyze unfolding of a protein and translocate the newly unfolded polypeptide chain through the axial channel of its hexameric ring structure during the protein disaggregation reaction. Consistently, recent cryoEM structures show unfolded model protein substrates residing in the axial channel of the hexameric ring structure of ClpB and Hsp104 [18, 28]. However, how these polypeptide chains arrive in the axial channel remains unknown. Is the substrate linearly translocated from one side of the hexameric ring, does the motor assemble around the substrate, or does the substrate enter through the open seam observed in many cryoEM structures of both ClpB and Hsp104 [28, 29]? Consequently, to address such questions, there is a need to develop, in solution, methods with the temporal resolution to complement these high spatial resolution structures.

For many reasons, direct measures of the translocation reaction catalyzed by the protein disaggregating machines has been difficult to acquire. Because of this, we have limited knowledge of the elementary steps in the protein unfolding and translocation reaction catalyzed by ClpB and Hsp104. In turn, this has limited our knowledge of the fundamental parameters describing the function, e.g. overall rates of unfolding and translocation, step-sizes (amino acids unfolded and/or translocated per turnover), ATP coupling efficiency, processivity, and directionality of translocation.

We recently reported the development and first application of a single turnover fluorescence stopped-flow approach that reports on the elementary kinetic steps in ClpB catalyzed protein unfolding of soluble model substrates [19]. Using that approach we reported quantitative information on the elementary kinetic steps describing enzyme catalyzed protein unfolding, the unfolding kinetic step-size, and processivity. Here we have employed that method to examine the ATP concentration dependence of the reaction and thereby the dependence of the elementary kinetic parameters on [ATP].

In our single turnover approach, it is essential that ClpB is pre-bound to the protein targeted for unfolding before initiating the unfolding reaction. The pre-binding reaction requires the use of non-hydrolysable or slowly hydrolysable nucleotide analogues because ClpB requires nucleoside triphosphate binding to assemble into hexameric rings with protein binding activity [13, 14, 23]. Curiously, we have shown that only ATPγS will serve the purpose of pre-assembly and binding, and the non-hydrolysable analogues like AMP-PNP and AMP-PCP will not substitute [16]. Thus, ATPγS serves a unique purpose that is not fully understood. Consequently, ATPγS is always present in the single-turnover experiments reported here.

Wickner and Coworkers showed that they could activate ClpB to catalyze protein disaggregation with a 1:1 mix of ATP:ATPγS that alleviated the need to include the co-chaperones, KJE, thereby simplifying *in vitro* studies [17]. In that work they showed that both the ATPase activity and the protein reactivation function could be optimized in the presence of a mixture of ATP and ATPγS. Both functions were shown to increase up to 1:1 (ATP:ATPγS) and then precipitously decreased at excess ATP.

In the single turnover stopped-flow experiments reported here, the presence of ATPγS in our experiments serves two purposes. First, as stated above, it is required to form an active hexamer with protein binding activity, and second, it alleviates the need for the c-chaperones, KJE. However, in contrast to what has been previously reported, we did not detect a decrease in the rate or elementary rate constant for protein unfolding at large excesses of ATP over ATPγS. Rather, we observe the expected saturable dependence of the rate on [ATP] that saturates at ∼50-fold excess ATP over ATPγS.

The rate of ClpB catalyzed protein unfolding depends cooperatively on ATP concentration at the fixed ATPγS = 150 μM. Although the cooperativity parameter or Hill coefficient is modest at ∼1.5, it does suggest cooperativity between ATP binding and hydrolysis sites. However, substantially more work is needed to determine if this is inter-or intra-subunit cooperativity or some combination of both. Nevertheless, with the development of this approach we are well positioned to begin addressing such questions.

ATPase studies on ClpB from *T. thermophilus* reported cooperativity with a hill coefficient of ∼2.4 and a *k*_*cat*_ ∼0.07 s^-1^ [30]. *S. cerevisiae* Hsp104 also exhibited cooperative ATP turnover with a Hill coefficient of ∼2.3 and *k*_*cat*_ ∼1.3 s^-1^ [31]. *E. coli* ClpB exhibited a similar Hill coefficient of ∼2.5 and *k*_*cat*_ ∼0.13 s^-1^. However, when *E. coli* ClpB was saturated with α-casein the Hill coefficient decreased to ∼1.4 and the *k*_*cat*_ increased to ∼0.3 s^-1^ monomer^-1^ [18]. Interestingly, the Hill coefficient of ∼1.4 determined in our analysis of the overall unfolding rate vs. ATP is identical to that reported by Deville et. al [18].

Bulk measurements of ATP turnover will reflect all oligomers hydrolyzing ATP in solution. We have shown that ClpB resides in a monomer, dimer, tetramer, hexamer equilibrium with a nucleotide binding stoichiometry of 1, 3, 7, and 12, respectively [13, 14, 23]. Thus, in a bulk ATPase experiment, a substantial amount of ATP hydrolysis may be coming from smaller order oligomers that are not catalyzing protein unfolding and translocation even in the presence of polypeptide substrate. Importantly, the ATP concentration dependence of the unfolding rate constant, reported here, is sensitive only to bound hexamers since only hexamers bind and unfold the protein substrates. Thus, the ATP concentration dependence of the observed rate constant only reflects the activity of hexamers that are catalyzing unfolding and translocation. Therefore, the unfolding rate constant reported here of *k*_*U*_ ∼ 0.34 s^-1^ is per hexamer. In contrast, the previous value of *k*_*cat*_ ∼ 0.3 s^-1^ is reported to be per monomer [18]. Those experiments were carried out at 0.5 μM ClpB. When accounting for the ligand-linked assembly equilibrium, a monomer concentration of 0.5 μM predicts a hexamer concentration of ∼ 1.5 nM ClpB_6_, which, in turn, predicts a *k*_*cat*_ > 100 s^-1^ per hexamer. The disparity between the two measurements likely represents a significant amount of uncoupled ATP hydrolysis detected in bulk ATP turnover.

Interestingly, from our single turnover stopped-flow experiments, the rate and elementary rate constant increases with increasing ATP and saturates at ∼50-fold excess ATP over ATPγS. In fact, both the elementary rate constant and the overall rate increased up to a maximum value with no suggestion of decreasing at substantial excess of ATP over ATPγS, as has been previously reported from other approaches [17]. This implies that once hexameric ClpB is loaded onto the protein substrate then protein unfolding will ensue with only ATP. Thus, the previously reported multiple turnover experiments reporting on substrate reactivation likely reflect the inability of ClpB to bind the substrate in the presence of large excesses of ATP over ATPγS. Thus, a likely interpretation is that the mixture of ATP and ATPγS is most important for stabilizing a bound complex but once ClpB catalyzed protein unfolding ensues, ATPγS is no longer needed. That is to say, slow ATP hydrolysis at one or more sites may not be required.

ATPγS stabilization of the bound complex vs. impacts on the underlying mechanism is supported by the observation that the peak amplitude decreases with increasing [ATP] over [ATPγS], shown in Figure 4 A and B. Under single turnover conditions, the peak amplitude is proportional to the amount of ClpB bound, 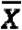, and the processivity raised to the power of the number of steps, *P*^*n*^. Consequently, the reduction in peak amplitude with increasing [ATP] is the result of either a reduced processivity or a reduction in the extent of binding.

The extent of binding is expected to be the same for the experiments with RepA-Titin_3_ as a function of [ATP]. This is because, in the first mixing event, the concentrations of ClpB, RepA-Titin_3_, ATPγS, and incubation time, Δt_1_ are all the same. Thus, if the extent of binding is different for each experiment carried out at different [ATP] then it must be the consequence of a perturbation to the bound complex that occurs on mixing with ATP, i.e. the second mixing event.

Upon rapid mixing with ATP the bound hexamers may disassemble because of the two-fold reduction in the ClpB and ATPγS concentrations. This interpretation is particularly attractive as ClpB has been reported to be in “rapid subunit exchange” [32]. However, from extensive examination of the assembly mechanism we have shown that the lifetime of the ClpB hexamer, at these ATPγS concentrations, is between 1,000 and 10,000 seconds [14]. Thus, we favor an interpretation where some fraction of ClpB either does not initiate unfolding and translocation or some fraction dissociates upon rapid mixing with ATP. In these experiments we cannot distinguish between these two possibilities as both explanations present a reduction in the extent of binding.

The alternate explanation to the reduction in peak amplitude as a function of increasing [ATP] is that the processivity is an ATP concentration dependent parameter. To test this hypothesis we quantified the processivity in two ways. One way is under the express assumption that processivity is independent of [ATP]. The second approach compares the peak amplitude as a function of substrate length, which assumes processivity depends on substrate length and not [ATP]. Determination of processivity under both assumptions led to a value of *P* ∼ 0.7. We take the agreement between the two strategies to suggest that processivity is independent of [ATP]. However, further refinement and development of our approach to quantify processivity is needed.

From the results presented in this manuscript we propose that, at saturating [ATP], ClpB induces the cooperative unfolding of a complete TitinI27 domain of 98 amino acids, which is represented by the kinetic step-size m ∼100 amino acids. This unfolding event is followed by rapid and undetected translocation up to the next folded domain. However, at sub-saturating [ATP], ClpB still induces cooperative unfolding of a complete TitinI27 domain but translocation becomes partially rate-limiting, which leads to an apparent reduced kinetic step-size as small as ∼ 50 amino acids.

Thus, the processivity of P ∼0.7 represents the processivity of protein unfolding. That is to say, the probability of unfolding vs. dissociation. Going forward, it will be of interest to determine the dependence of processivity on the thermodynamic stability of a variety of different protein substrates and even the directionality of translocation.

## Materials and Methods

### Buffers and reagents

Reagent-grade chemicals were used to prepare buffers using 18 MΩ deionized water from a Purelab Ultra Genetic system (Evoqua, Warrendale, PA). All reagents were dialyzed into buffer H200, which contains 25 mM HEPES, pH = 7.5 at 25 °C, 10 mM MgCl_2_, 200 mM NaCl, 2 mM 2-mercaptoethanol and 10 % (v/v) Glycerol. *E. coli* ClpB was purified as described [33]. All ClpB concentrations are reported in monomer units. ATP and ATPgS were purchased from Thermo Fischer Scientific (Waltham, MA) and CalBiochem (La Jolla, CA), respectively. Both ATP and ATPgS were dialyzed into H200 using 100-500 Da molecular weight cut-off dialysis tubing (Thermo Fischer Scientific, Waltham, MA). a-casein was purchased from Sigma-Aldrich (Darmstadt, Germany), dissolved in 6 M Guanidine hydrochloride, 20 mM HEPES, pH 7 at 25 °C, and dialyzed into H200 using 10 kDa molecular weight cut-off dialysis tubing (Thermo Fischer Scientific, Waltham, MA). Purification and sample preparation of RepA-Titin_X_ were carried out as described in [22].

### Structures of RepA-Titin_x_

The 70 amino acids of the Phage P1 RepA protein (RepA_70_) are present on the substrate’s N terminus, which is displayed as unstructured regions on all substrates (see Fig. 1 A). RepA_70_ is followed by the folded b sandwich structure of TitinI27. On the C-terminus of each substrate, Cysteine is shown as space-filling to which Alexa Fluor 555 is covalently attached (not shown). The difference between the three substrates is in the number of tandem repeats of TitinI27, which are connected by unstructured regions. For RepA-Titin_1_, there is only 1 repeat of TitinI27 present, and so on.

### Sequential mix stopped-flow experiments

Sequential mix mode on SX20 Applied Photophysics stopped-flow fluorometer (Leatherhead, U. K.) was used to conduct experiments at 25°C in H200 buffer. ClpB and RepA-Titin_X_ were dialyzed into the H200 buffer using dialysis tubing of 50 kDa and 10 kDa molecular weight cutoff, respectively.

#### [ClpB] dependence experiments schematized in Fig. 1 C

Varying concentrations of ClpB monomer (4 μM, 8 μM, and 14 μM) are rapidly mixed with 600 μM [ATPγS] and 200 nM [RepA-Titin_X_]. After mixing, they travel to the ageing loop where the reagents age for fixed Δt_1_ = 10 min. In this aging period, we expect the formation of ClpB_6_ in the presence of ATPγS and the binding of ClpB_6_ with RepA-Titin_X_ to form a pre-bound complex. After Δt_1_ has occurred, the pre-bound complex is rapidly mixed with 1 mM ATP and 40 μM α-casein in the mixing chamber 2. Then, they travel together to the observation chamber where fluorescence is monitored by exciting Alexa Fluor 555 at 520 nm. The emission signal is collected using a long pass filter of 570 nm. The reagents in syringes 1 and 2 are diluted four-fold, whereas those in syringe 3 are diluted two-fold. For consistency in this paper, we refer to the final concentrations of all reagents. For example, 2 μM ClpB in the observation channel results from 8 μM ClpB in syringe 1.

#### [ATP] dependence experiments schematized in Fig. 2 B

Similar experiments were carried out as described in [ClpB] dependence experiments except using a fixed [ClpB] = 2 μM and varying [ATP].

Processing of raw time-courses was carried out by using Eq 6.

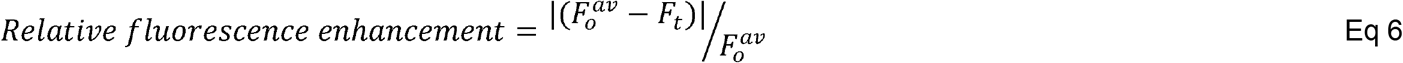

where 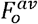 is the average of the first few constant raw fluorescence data points and *F*_*t*_ is raw fluorescence signal at a given time.

#### Fitting experimental time-courses to Scheme 1

We plot peak time versus the total substrate length for each set of experimental time courses at a particular condition, see Fig. 1 D and 2 C. The intercepts that we get from these plots are termed *C*. Reduced length, *L’*, is determined by subtracting *C* from the total length of the substrate, *L*. See Figs. 1 F and 2 E for average *C* values of [ClpB] and [ATP] dependence time-courses. We use *L’* in our fitting routine to exclude the substrate length unavailable for translocation and unfolding to ClpB, i.e., pre-translocated distance with ATPγS, dangle distance, and occluded length. Note that we determine the *C* value for each data set individually.

The custom-built MATLAB (MathWorks, Natick, MA) toolbox, MENOTR [34] was used to fit the time-courses to Scheme 1 (Fig. 3 A). The data sets at different [ClpB] and [ATP] were described using eq 7. For [ClpB] dependence data, *n* was replaced by *L’*/*m*, whereas for [ATP] dependence data, *n* was replaced by *L’*/*m* + *b*. Out of all parameters, *b*, m, k_U_, and k_end_ are kept global.

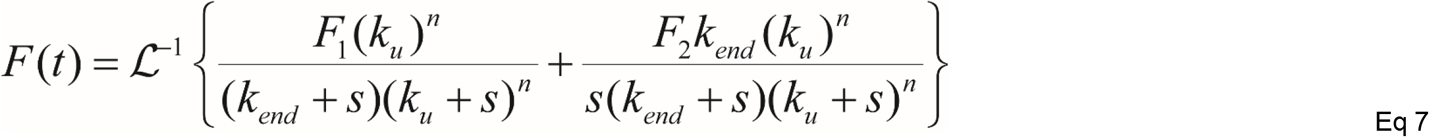

Where *F*_*1*_ and *F*_*2*_ are two fluorescence amplitudes for the last intermediate, I_(L-nm)_ and unfolded RepA-Titin_x_ respectively, *s* is the Laplace variable and other kinetic parameters are defined as per scheme 1, see Fig. 3 A.

#### Calculation of processivity

The fitting of time-courses was carried out as mentioned earlier, except that *n* was not replaced with *L’/m* + *b* instead all three lengths at a particular [ATP] were fit globally to a local *n* in eq 7. The *n* determined for RepA-Titin_3_ was used as an independent variable in Fig. 4 C. To note that processivity was determined for each replicate individually by investigating the peak amplitudes as function of number of steps.

## Supplemental Figures

**Supplemental Figure 1:**
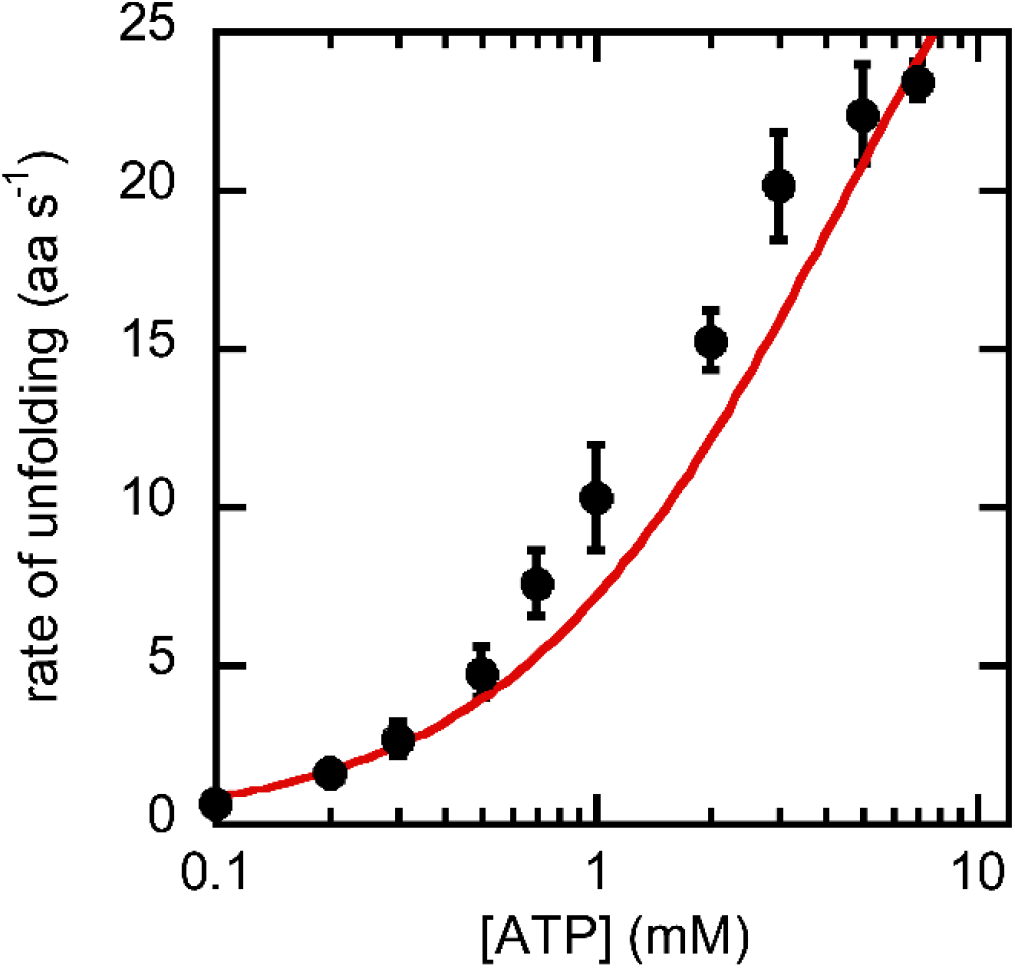
Plot of the rate of unfolding as a function of [ATP] fit to hyperbolic dependence described in eq 1 except *h* is fixed to 1.

**Supplemental Figure 2:**
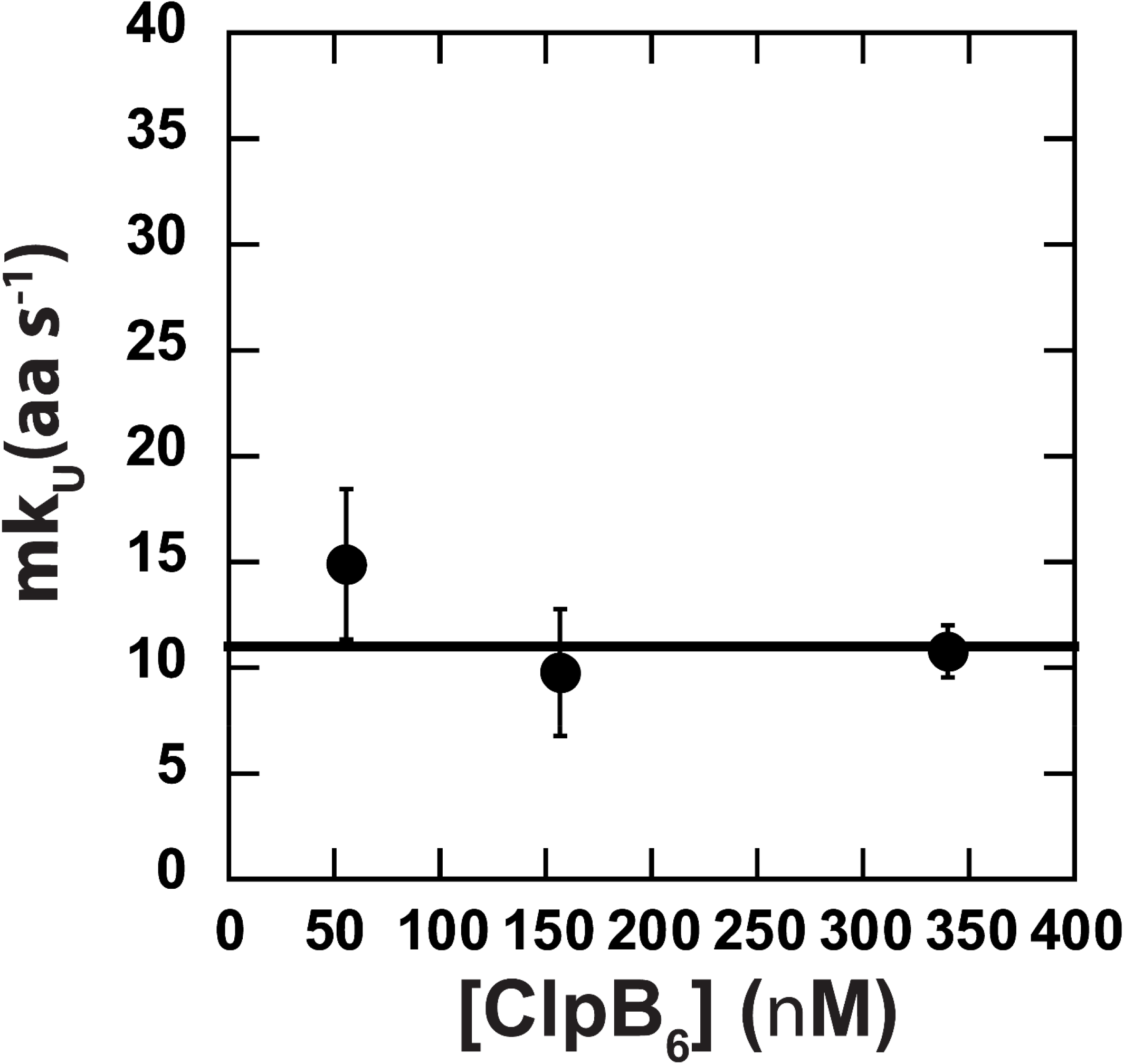
Fitting [ClpB] dependence time courses (see Fig. 3 A) to Scheme 1 in Fig. 2 A.The product of obtained parameters *m* and *k*_*U*_ is plotted as a function of [ClpB_6_]. This plot is fit to a constant at *m* = (11 ± 1) aa s^-1^. The error bars represented here are propagated from fitting errors of *m* and *k*_*U*_.

## Acknowledgment

We thank Kaila Fuller and members of the Lucius lab for their critical discussions of the results and the manuscript. This work was supported by the National Science Foundation (grant NSF MCB-1412624 to A.L.L.). Computational work was performed using the UAB High Performance Computing (HPC) Cheaha, which is supported in part by the National Science Foundation under Grants No. OAC-1541310, the University of Alabama at Birmingham, and the Alabama Innovation Fund.

